# A Marfan-Associated FBN1 Nonsense Mutation Mouse Model Reveals Adventitial Inflammation During Aneurysm Progression

**DOI:** 10.1101/2025.09.23.678101

**Authors:** Shichao Wu, Jiawei Zhao, Alejandro Ponce, Lucynda Pham, Daniel Xie, Donghong Ju, Shuping Yin, Francis Hernandez, Sean Jones, Clair Li, Charles Chung, Dragana Komnenov, Noreen F. Rossi, Dogukan Mizrak, Bo Yang, Zhe Yang, Maozhou Yang, Hui Li, Youming Xie, Kang Chen, Kezhong Zhang, Li Li

## Abstract

Marfan syndrome (MFS) is a genetic disorder caused by mutations in *fibrillin-1(FBN1)*, which encodes FBN1, a key structural component of the extracellular matrix. Mutations in *FBN1* influence the severity of aortic disease and therapeutic responses, with aortic aneurysm being the leading cause of mortality in patients with MFS. To investigate the mechanisms driving aneurysm progression, we generated a mouse model (*Fbn1^Q2469X/+^*) carrying the FBN1Q2467X nonsense mutation identified in MFS patients. This mutation results in FBN1 deficiency. Although *Fbn1^Q2469X/+^* mice appear normal, showing only mild, nonprogressive dilation of the aortic root and ascending aorta with minor reductions in blood pressure, homozygous *Fbn1^Q2469X/Q2469X^* mice develop spontaneous thoracic aortic aneurysms (TAA) that progress to rupture between 10 and 25 days of age, with 100% penetrance. Histopathology shows progressive vessel wall degeneration characterized by disorganized vascular smooth muscle cells, collagen loss, and elastic fiber fragmentation from early to late stages. RNA-seq analysis identifies inflammation as the dominant process in late-stage aneurysms. Immunofluorescence assay reveals inflammatory cells prominently localized to the adventitia near rupture sites, linking adventitial inflammation to aneurysm progression. This genetically modified *Fbn1^Q2469X/Q2469X^* mouse model consistently develops progressive aortic aneurysms and provides a reliable, cost-effective platform to investigate the molecular mechanisms of aneurysm progression and to evaluate therapeutic strategies in aneurysm diseases, including Marfan syndrome and related disorders.

## Introduction

Marfan syndrome (MFS) is a heritable connective tissue disorder that affects the cardiovascular, skeletal, ocular, and pulmonary systems^1–5^. Aortic aneurysm and dissection are the leading causes of mortality in MFS patients, underscoring the urgent need for improved understanding of disease mechanisms and therapeutic strategies. MFS is primarily caused by pathogenic variants in *FBN1*, which encodes fibrillin-1, a multifunctional extracellular matrix glycoprotein that regulates critical signaling pathways, including TGF-β, integrins, and growth factor bioavailability^4,6–10^. More than 3,000 FBN1 variants have been reported, including missense, nonsense, and frameshift mutations, each exerting distinct effects on fibrillin-1 structure, stability, or abundance; approximately one-third of MFS cases arise from *de novo* mutations without family history^4,7^. Clinical studies demonstrate that aneurysm severity varies depending on mutation class ^11–13^, suggesting that different molecular mechanisms converge on overlapping but distinct pathogenic pathways. Patients with haploinsufficiency (loss-of-function) variants such as nonsense or frameshift mutations typically develop more aggressive aortic disease than those with most dominant-negative missense variants, except for mutations in the neonatal FBN1 region (exon 25-33) as well as in the region of exons 26-49^12–14^. Understanding why reduced fibrillin-1 dosage leads to more severe vascular phenotypes remains an important unresolved question.

FBN1 mutations appear to influence therapeutic response in MFS patients. Angiotensin II receptor blockade with losartan has been shown to attenuate aortic root dilation in patients carrying nonsense and frameshift mutations, but its benefit is less pronounced in those with missense variants^15–18^. These findings suggest that molecular context (haploinsufficiency versus dominant-negative mutation) determines not only disease severity but also treatment efficacy. The development of mutation-relevant animal models is therefore essential for understanding the underlying mechanisms of aortic disease and for guiding precision therapies in MFS.

To address this gap, we used the CRISPR/Cas9 genome editing technology to generate a new Marfan mouse model *Fbn1^Q2469X/+^*, harboring the FBN1Q2467X nonsense mutation found in Marfan patients^21,22^. Heterozygous mice develop slightly increase in the aortic root and the ascending aorta, while homozygous mice uniformly develop TAA that rupture within 10–25 days of age. Histopathological and transcriptomic analyses reveal smooth muscle cell remodeling, extracellular matrix degradation, and oxidative stress, with inflammatory cells prominently localized in the adventitia near rupture sites. These findings identify adventitial inflammation as a significant feature of late-stage aneurysms.

## Material and Methods

### Generation of *Fbn1^Q2469X/+^* Mice

The FBN1 gene is highly conserved between humans and mice: the human nonsense mutation *FBN1* c.7399C>T (p.Q2467X) in exon 59 corresponds to mouse *Fbn1* c.7405C>T (p.Q2469X) in exon 60 (Figure 1 and Figure 2A). The mouse *Fbn1* gene (GenBank accession number NM_007993.2; Ensembl ENSMUSG00000027204) is located on chromosome 2 and contains 66 exons, with the ATG start codon in exon 2 and the TAA stop codon in exon 66.

**Figure 1.**
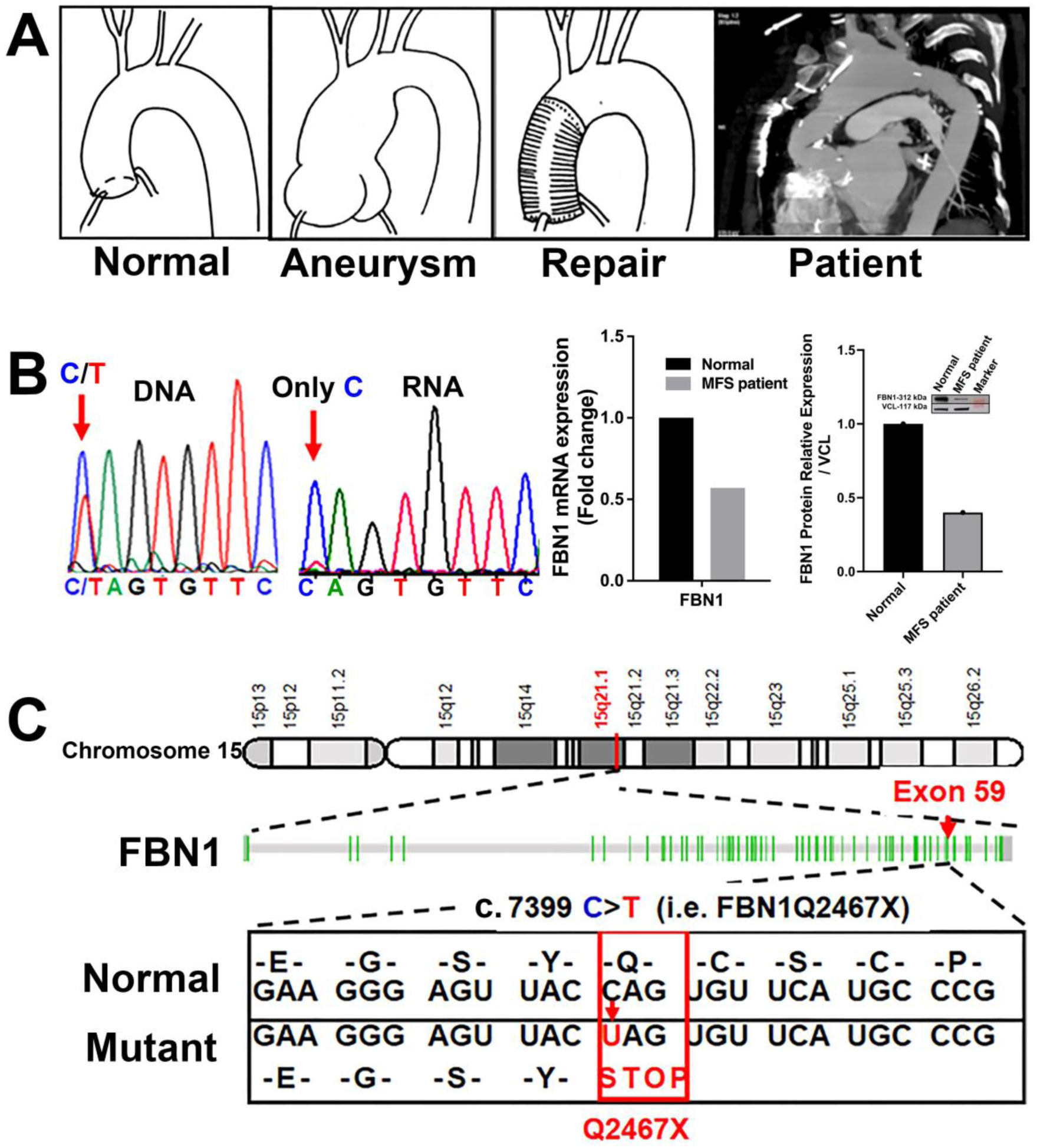
The *FBN1Q2467X* nonsense mutation associated with Marfan syndrome. (A) CT scan of a Marfan syndrome patient carrying the *FBN1Q2467X* mutation shows aneurysms in the thoracic aortic arch and evidence of prior surgical repair of the aortic root and ascending aorta.(B) Sanger sequencing chromatograms from the patient show both C and T peaks at position c.7399 in genomic DNA, whereas only the C allele is detected in cDNA, indicating degradation of transcripts carrying the mutant T allele. qPCR and Western blot analysis demonstrate ∼50% reduced expression of *FBN1* mRNA and protein in the patient compared with a non-Marfan control. (C) Schematic representation of *FBN1* on chromosome 15, which spans 66 exons. The c.7399C>T transition in exon 59 introduces a premature stop codon (UAG), resulting in a nonsense mutation (Q2467X).

**Figure 2.**
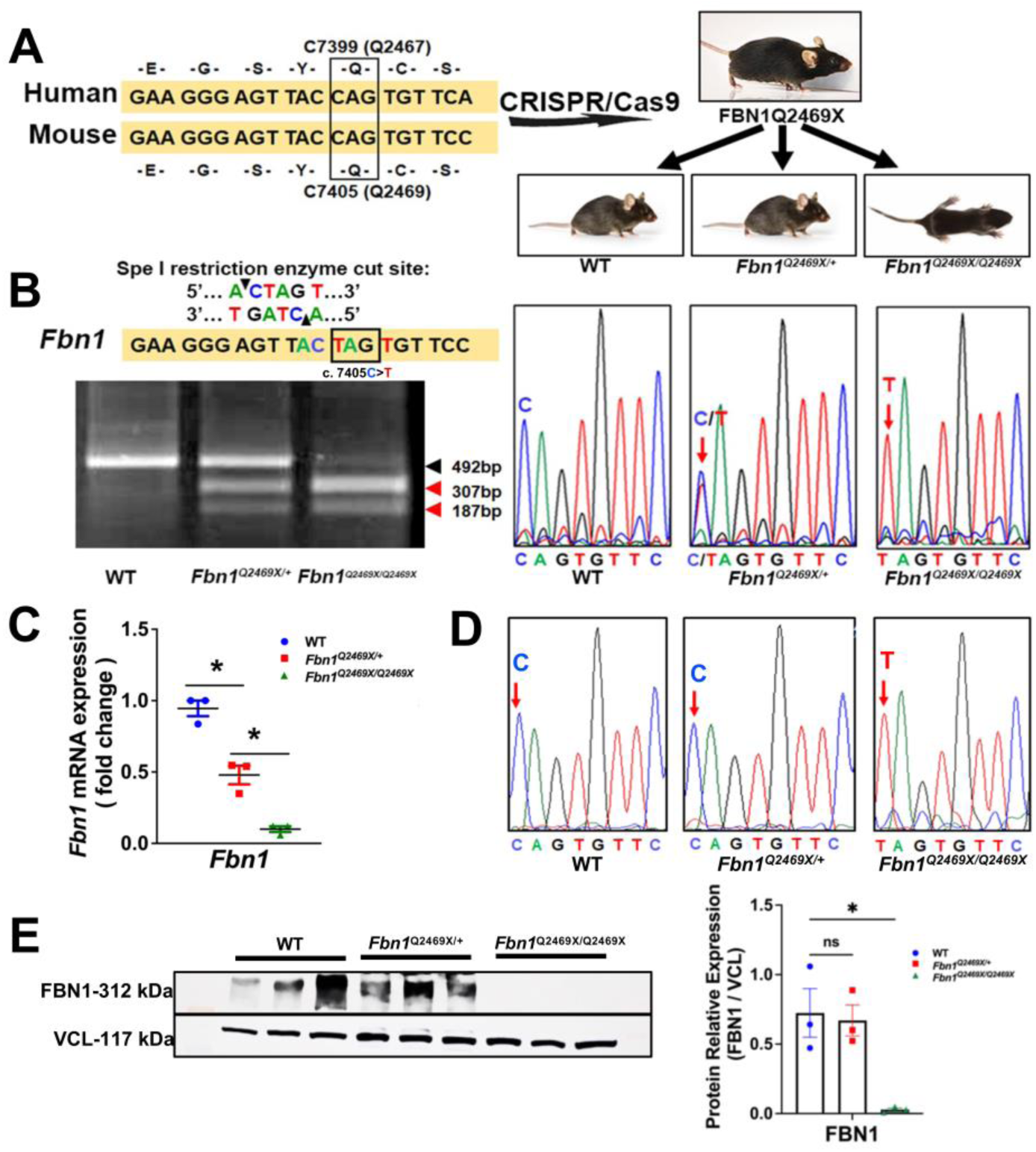
Generation and validation of the FBN1Q2469X Marfan mouse model. (A) Conservation of FBN1 between human and mouse. The human *c.7399C>T* mutation (Q2467X) in exon 59 corresponds to mouse *c.7405C>T* (Q2469X) in exon 60. CRISPR/Cas9 genome editing was used to introduce the *c.7405C>T* nonsense mutation into the mouse *Fbn1* gene, generating FBN1Q2469X mice. Breeding produced wild-type (*Fbn1^+/+^),* heterozygous (*Fbn1^Q2469X/+^*) and homozygous (*Fbn1^Q2469X/Q2469X^*) genotypes. Representative pups at 10 days of age are shown. (B) Genotyping strategy. PCR products encompassing the mutation site were digested with SpeI, which is reconstituted only in the mutant allele (ACTAGT). The wild-type allele yields a single 492 bp band, while the mutant allele produces 307 bp and 187 bp fragments. Sanger sequencing chromatograms confirm the presence of the *c.7405C>T* mutation in heterozygous and homozygous mice. (C) *Fbn1* mRNA expression. qPCR shows a ∼50% reduction of *Fbn1* mRNA in heterozygotes and >90% reduction in homozygotes (n=3; one-way ANOVA, *P*<0.05). (D) cDNA sequence analysis. Agarose gel electrophoresis of RT-PCR products (left) and corresponding Sanger chromatograms (right) demonstrate selective loss of the mutant transcript in heterozygotes, consistent with nonsense-mediated decay, while the mutant allele is retained in homozygotes. (E) FBN1 protein expression. Western blotting shows slightly reduced fibrillin-1 protein in heterozygotes and near-complete loss in homozygotes, relative to wild type. Vinculin (VCL) was used as a loading control (n=3; *P*<0.05).

The *Fbn1^Q2469X^*allele was generated using CRISPR/Cas9-mediated genome editing (Cyagen Biosciences, Santa Clara, CA). Cas9 mRNA, sgRNA, and a donor oligonucleotide carrying the Q2469X mutation (CAG>TAG) were co-injected into zygotes. Homology-directed repair introduced the nonsense mutation into exon 60, and a silent mutation (TCC>TCG) was incorporated to prevent re-cutting by sgRNA. Pups were genotyped by PCR and confirmed by Sanger sequencing. Positive founders were bred to establish germline transmission. Sequence analysis of five predicted off-target sites showed no evidence of unintended mutations. Wild-type DNA was sequenced in parallel as a negative control to exclude artifacts from sequencing chromatograms.

The resulting *Fbn1^Q2469X/+^* line was designated as *Fbn1^tmQ2469XWSUlili^* (abbreviated *Fbn1^Q2469X/+^* or FBN1Q2469X when deposited at The Jackson Laboratory), following the official guidelines for mouse strain nomenclature (http://www.informatics.jax.org/mgihome/nomen/gene.shtml).

All animal procedures were approved by the Institutional Animal Care and Use Committee (IACUC) of Wayne State University (WSU) and conformed to the NIH Guide for the Care and Use of Laboratory Animals. The *Fbn1^Q2469X/+^* mice were maintained on a C57BL/6J background, and both sexes were used in experiments.

### Mouse Genotyping

Genomic DNA was isolated from pup toes or adult ear punches using DirectPCR lysis buffer (Viagen Biotech) supplemented with Proteinase K (10 mg/mL). Samples were digested overnight at 55°C, followed by heat inactivation at 98 °C for 10 minutes. PCR amplification was performed using REDTaq ReadyMix (Sigma-Aldrich) with the following primers: forward 5′-AACAGGGACCGCCTGTGTAGGTAG-3′ and reverse 5′-GCTCAGGGTTATCCTCAGGGACC-3′, yielding a 492 bp product. PCR conditions were: initial denaturation at 95 °C for 1 minute; 35 cycles of denaturation at 95 °C for 10 seconds, annealing at 62 °C for 30 seconds, and extension at 72 °C for 30 seconds; and a final extension at 72 °C for 10 minutes.

The *Fbn1^Q2469X/+^* allele introduces a SpeI restriction site (ACTAGT), allowing differentiation of wild-type (492 bp), *Fbn1^Q2469X/+^* heterozygous (492 bp, 305 bp and 187 bp), and *Fbn1^Q2469X/Q2469X^* homozygous (305 bp and 187 bp) genotypes. PCR products were digested with SpeI (New England Biolabs) in a 15 μL reaction containing 1.5 μL CutSmart buffer, 3 \μL ddH₂O, and 0.5 μL SpeI enzyme per reaction. Five microliters of the digestion mix was added to 10 μL PCR product and incubated at 37 °C for 3 hours. Digested fragments were resolved by a 1.5% agarose gel in TBE buffer at 100 V for 30 minutes and visualized by ethidium bromide staining (Figure 2B).

### Sequence Verification of the *Fbn1* Mutation

The presence of the FBN1 mutation was verified by Sanger sequencing (GENEWIZ). PCR products amplified with primers flanking the Q2467X site (Table 1) were submitted together with 40 μl of 5 μM forward and reverse primers. Forward and reverse reads were aligned to the *FBN1* reference sequence using NCBI BLAST (nucleotide database). Sequences were examined for the wild-type allele (CATTGTTCC) or the mutant allele (TAGTGTTCG). Sequence trace files were visualized using Chromas Lite software to generate chromatograms for DNA and cDNA. Representative chromatograms are shown for a Marfan patient carrying the FBN1Q2467X mutation and for mice of three genotypes (*Fbn1^+/+^, Fbn1^Q2469X/+^*and *Fbn1^Q2469X/Q2469X^*)(Figure 1B, Figure 2B, 2D).

**Table 1.**
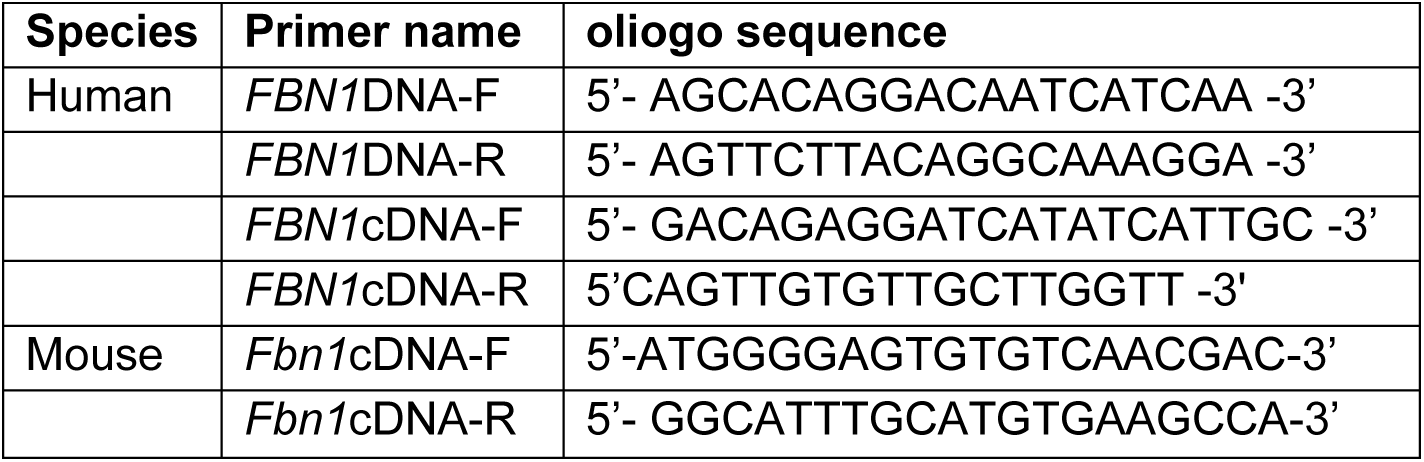
Primers used to sequence *FBN1* gene and cDNA.

### Blood Pressure Measurement

Systolic blood pressure was measured in both *Fbn1^+/+^*and *Fbn1^Q2469X/+^* mice at 3 months of age using the CODA™ Non-Invasive Blood Pressure System (Kent Scientific, Torrington, CT). Mice underwent a 10-day testing protocol, with a 2-day rest period between two 5-day sessions. The first 5 days were used to acclimate the animals to the holders and equipment, and the subsequent 5 days were used for data collection. Blood pressure values were averaged across the final 5-day recording period. Equipment performance and signal quality were monitored continuously, and measurements were repeated if artifacts or cuff malfunctions were detected.

### Echocardiography

Adult *Fbn1^+/+^* and *Fbn1^Q2469/+^* mice were assessed by transthoracic echocardiography using the Vevo 3100 system equipped with an MS550 transducer, as previously described^23^. Mice were anesthetized with 3% isoflurane in oxygen for induction and maintained at 1.5–2% isoflurane during imaging. Animals were positioned supine on a heated platform, and aortic dimensions were measured at the aortic root and ascending aorta. Vessel mechanical properties were derived from echocardiographic recordings.

### Latex Injection for Aneurysm Visualization

To visualize aortic morphology during aneurysm progression, vascular casting with yellow latex was performed as previously described^24^ with modifications. After euthanasia, the abdominal and thoracic cavities were opened to expose the heart and major vessels. The right atrium was incised following perfusion with PBS through the left ventricle. To preserve native aortic morphology, PBS was replaced with 4% neutral-buffered formalin for fixation. Two milliliters of yellow latex medium (Ward’s Science, catalog #470024-616) were then slowly injected through the left ventricle. The left ventricle and right atrium were clamped with Serrefine clamps, and mice were immersed in 4% neutral-buffered formalin overnight at 4 °C. The following day, formalin was replaced with 70% ethanol for ∼2 hours at 4 °C before dissection and imaging.

### Aorta Isolation and Processing for RNA-seq, Western Blotting, and Histology

*Fbn1^+/+^* and *Fbn1^Q2469X/Q2469X^*mice at 13–15 days of age were euthanized, and the thoracic cavity was opened. The right atrium was incised to allow blood drainage, followed by perfusion with ice-cold PBS through the left ventricle until the effluent was clear and the liver appeared pale. Perfusion was then continued with 4% paraformaldehyde (PFA) to fix the aorta *in situ*. Hearts and aortas were collected, post-fixed in 4% PFA for 8–12 hours at 4 °C, and cryoprotected in 30% sucrose at 4 °C overnight or until tissues reached equilibrium (floating midway in solution). Tissues were embedded in OCT compound on dry ice and stored at −80 °C. Cryosections were cut at 5 *μm* thickness and mounted onto glass slides, which were stored at −20 °C until use.

### Histology analyses

OCT-embedded aortic sections (5 *μm*) were used for hematoxylin and eosin (H&E), collagen (Picro Sirius Red Stain Kit, Abcam, Cat. #ab150681), and elastic fiber staining (Verhoeff–Van Gieson Elastin Kit, Polysciences, Cat. #25089). Prior to staining, three preparatory steps were performed: (1) slides were removed from the −20 °C freezer and allowed to equilibrate at room temperature for 30 minutes; (2) sections were fixed in cold methanol:acetone (1:1) for 5 minutes; and (3) slides were rinsed twice in PBS (3 minutes each). Stained sections were imaged by light microscopy using a Leica DM4000B microscope and LAS software at 400× magnification. Multiple images were acquired and stitched digitally to generate high-resolution composite images.

### Immunofluorescence

OCT-embedded aortic sections (5 *μm)* were air-dried for 30 minutes at room temperature, fixed in cold acetone:methanol (1:1) for 10 minutes, and rinsed in PBS. Sections were permeabilized with 0.1% Triton X-100 in PBS (2 × 5 minutes), followed by washing in PBST (0.1% Tween-20 in PBS). Endogenous peroxidase activity was quenched with 3% H₂O₂ in methanol (10 minutes), and sections were rinsed again in PBST (3 × 5 minutes). Tissue boundaries were circled with a hydrophobic barrier pen, and sections were blocked with PBS containing 10% serum (matched to the host species of the secondary antibody) and 1% BSA for 1 hour. Slides were incubated overnight at 4 °C with primary antibodies diluted in 50% blocking buffer. After washing (3 × 5 minutes, PBST), sections were incubated with secondary antibodies (diluted in 50% blocking buffer) for 2 hours at room temperature, followed by additional washes (5 × 5 minutes, PBST). Sections were dehydrated in graded ethanol (95% for 20 seconds, 100% twice for 30 seconds each), cleared in xylene (3 × 1 minute), and mounted with DAPI-containing medium (Vector Laboratories, Cat. #H-2000). Fluorescence images were captured using a Leica DM4000B microscope with LAS software at 400× magnification. Multiple fields were acquired per section. All antibodies were tested in triplicate, and negative controls omitting the primary antibody were included in each experiment.

### RNA extraction for RNA-seq and qPCR

Thoracic aorta from *Fbn1^+/+^*, *Fbn1^Q2469X/+^* and *Fbn1^Q2469X/Q2469X^* mice at 13 day old pups (n=3) were isolated and preserved in RNAlater and store in –70^0^C. RNA extraction, library preparation, RNA-seq and transcriptome analysis was performed by Admera Health Inc. (New Jersey, United States): sample were isolated using Qiagen RNeasy Plus Micro Kit (Qiagen, Hilden, Germany) with a modified protocol. RNA sample quality was assessed by Agilent RNA 6000 Pico Reagent on Agilent 2100 Bioanalyzer (Agilent Technologies, Santa Clara, CA) and quantified by Qubit RNA HS assay (ThermoFisher, Waltham, MA). Ribosomal RNA depletion and library preparation was performed for each sample with TruSeq Stranded Total RNA Library Prep Gold (Illumina Inc., San Diego, CA), following manufacturer’s instructions. Average final library size is about 300 bp. Illumina 8-nt dual-indices were used for multiplexing. Samples were pooled and sequenced on Illumina HiSeq X sequencer for 150 bp read length in paired-end mode, with an output of 80 million reads per sample (40M reads in each direction).

### RNA extraction from aorta tissues (flash freeze in liquid N2 and stored at –70^0^C)

cDNA for qPCR was synthesized using Superscript III reverse transcriptase (Invitrogen). Real-time PCR was performed using the Step-One Plus system (Applied Biosystems) in the presence of SYBR Green. *Gapdh* and *U6* was used as internal control. Changes in mRNA expression were expressed as fold change relative to internal control. All gene specific PCR primers were designed to flank at least 2 exons.

### RNA-seq Analyses

RNA isolation and sequencing was performed by Admera Health Inc. Quality of raw reads was assessed using FastQC (v0.11.8). Adapter sequences and low-quality bases were trimmed with Trimmomatic (v0.38). Clean reads were aligned to the mouse reference genome (GRCm38/mm10) using STAR Aligner (v2.7.1a). Gene-level reading counts were obtained with HTSeq-count (v0.11.2). Mapping quality was evaluated with Picard tools (v2.20.4; https://broadinstitute.github.io/picard/). Transcript abundances were estimated as fragments per kilobase per million mapped reads (FPKM) using StringTie (v2.0.4). Differential gene expression analysis was performed with DESeq2. Raw count data were normalized using the variance stabilizing transformation (VST) implemented in DESeq2 to obtain homoscedastic, log2-scale expression values. The top 2,000 most variable genes across samples were selected for heatmap visualization. For each gene, expression values were mean-centered to generate z-scores, which determined the heatmap color gradient. VST-normalized values were also used to compute pairwise Pearson correlations between samples. Sample distance heatmaps were constructed using the distance metric √(1 − r²), where *r* is the Pearson correlation coefficient.

### Protein Extraction and Western Blotting

Thoracic aortas were collected from 13-day-old *Fbn1^+/+^ Fbn1^Q2469X/+^* and *Fbn1^Q2469X/Q2469X^*mice (n = 3 per group). Protein lysates were prepared, and concentrations were determined using the BCA assay (Thermo Fisher Scientific). Equal amounts of protein were separated on 4–12% Bis–Tris NuPAGE Mini Gels (Invitrogen) and transferred onto PVDF membranes (Millipore) using the XCell II Blot Module (Invitrogen). A Spectra™ Multicolor Broad Range Protein Ladder (Thermo Fisher Scientific, Cat. #26634) was included for molecular weight estimation. Membranes were blocked with 5% nonfat milk at room temperature for 1 hour and incubated overnight at 4 °C with primary antibodies against fibrillin-1 (Abcam, Cat. #ab231094, 0.2 μg/mL; ∼312 kDa) and vinculin (Santa Cruz, Cat. #sc73614, 1:100; ∼117 kDa). After washing, membranes were incubated with HRP-conjugated secondary antibodies (anti-rabbit IgG, Abcam, Cat. #ab6721, 1:2000; anti-mouse IgG, Sigma-Aldrich, Cat. #AP130, 1:3000) for 1.5 hours at room temperature. Signals were detected by enhanced chemiluminescence (SuperSignal West Pico Chemiluminescent Substrate, Thermo Fisher Scientific, Cat. #21050) and imaged using the KwikQuant Imager (Kindle Biosciences).

### Statistical analyses

To Observed genotype frequencies were compared with expected Mendelian ratios using the chi-square test. Survival of *Fbn1^Q2469X/Q2469X^* mice was assessed by Kaplan–Meier analysis, and survival curves were compared using the log-rank test. Group comparisons were performed using two-tailed Student’s *t* tests. All statistical analyses were conducted with GraphPad Prism 9 (GraphPad Software, San Diego, CA). Data are presented as mean ± SEM, and the number of experimental replicates is provided in the figure legends. A *P* value < 0.05 was considered statistically significant.

## Results

### The FBN1Q2467X nonsense mutation in Marfan patients is associated with reduced FBN1 expression

Typically, MFS patients first present with aneurysms at the aortic root and ascending aorta that require surgical repair^5^. In the case shown here, a MFS patient with the *FBN1* c.7399C>T (p.Q2467X) mutation developed additional aneurysms in the aortic arch following repair of the aortic root and ascending aorta (Figure 1A). Sequence chromatogram analysis of DNA and RNA isolated from aneurysm tissue confirmed the presence of the mutation. Genomic DNA revealed heterozygosity at nucleotide position c.7399 (C/T), whereas cDNA from the same tissue showed only the wild-type C allele, indicating selective degradation of the mutant transcript by nonsense-mediated decay (Figure 1B). Consistent with this, qPCR and Western blot analyses demonstrated ∼50% reductions in *FBN1* mRNA and protein expression in the patient sample compared with a non-Marfan control.

The *FBN1* gene is located on chromosome 15q21.1 and comprises 66 exons encoding a 9,847 bp mRNA and a fibrillin-1 protein of 2,873 amino acids (∼350 kDa) (Table S1). Genetic testing of MFS patients has identified a c.7399C>T transition in exon 59, which converts the CAG codon for glutamine (Q2467) into a premature stop codon (TAG), generating the *FBN1Q2467X* nonsense mutation^21,22^. This mutation lies within the calcium-binding epidermal growth factor–like domain 39 (Figure 1C).

### CRISPR/Cas9-mediated generation of a Marfan mouse model carrying the orthologous FBN1Q2469X mutation

*FBN1* gene is highly conserved between human and mouse with 96% conservation in protein identity and 98% in protein similarity (Table S1). To model the human *FBN1Q2467X* nonsense mutation, we used CRISPR/Cas9 genome editing to introduce the orthologous mutation (*c.7405C>T,* Q2469X) into the mouse *Fbn1* gene (Figure 2A). The resulting *Fbn1*^Q2469X/+^ mice bred to produce *Fbn1^+/+^*, *Fbn1^Q2469X/+^* and *Fbn1^Q2469X/Q2469X^* mouse at Mendelian ratio. Although *Fbn1^Q2469X/Q2469X^*mouse is indistinguishable from the littermates at birth, some *Fbn1^Q2469X/Q2469X^* mice were runted 10 days after birth and displayed early lethality (Figure. 2A).

Since the C to T mutation creates a Spe1 restriction site that is only present in the *Fbn1* mutant allele. Genotyping was confirmed by PCR amplification of the target locus followed by SpeI digestion, which distinguishes the wild-type allele (492 bp) from the mutant allele (307 and 187 bp fragments). Sanger sequencing further validated the presence of the *c.7405C>T* mutation in heterozygous and homozygous mice (Figure 2B).

A nonsense mutation in *Fbn1* could either generate a truncated fibrillin-1 protein or trigger selective degradation of mutant transcripts via nonsense-mediated decay (NMD), resulting in FBN1 deficiency. To distinguish between these possibilities, we assessed *Fbn1* expression in mouse aortas, qPCR revealed that *Fbn1* mRNA levels were reduced by ∼50% in heterozygotes and by ∼95% in homozygotes compared with wild-type littermates (Figure 2C). Sequencing of RT-PCR products demonstrated that only the wild-type C nucleotide, and not the mutant T, was detected in heterozygotes, indicating loss of mutant transcripts (Figure 2D). The substantial reduction (∼ 90%) of *Fbn1* mRNA and the absence of detectable fibrillin-1 protein in homozygotes further support NMD-mediated transcript degradation. These findings confirm that the *Fbn1Q2469X* nonsense mutation leads to FBN1 deficiency, rather than the production of truncated protein, in both heterozygous and homozygous mice.

Western blotting demonstrated a gene dosage–dependent reduction of FBN1 protein: modestly decreased in heterozygotes and severely decreased in homozygotes compared with wild-type controls. Vinculin was used as a loading control (Figure 2E). Together, these data establish that the Fbn1Q2469X allele induces haploinsufficiency through nonsense-mediated decay: providing a genetic mouse aneurysm model of the human FBN1Q2467X mutation.

### *Fbn1^Q2469X/+^* mice appear normal with non-significant aortic dilation and normal histology

*Fbn1^Q2469X/+^*mice were grossly indistinguishable from their wild-type littermates, showing no evidence of aortic dilation, rupture, or other overt cardiovascular abnormalities during the first year of life. Echocardiography revealed a mild but non-significant increase in the diameters of the aortic root (L1–L3) and ascending aorta in *Fbn1^Q2469X/+^*mice (Figure 3A-B). Blood-pressure measurements showed a reduction in both systolic and diastolic pressures in *Fbn1^Q2469X/+^* mice, but these differences did not reach statistical significance (systolic *P* = 0.066; diastolic *P* = 0.4831; Figure 3C). Histological examination of the aortic wall revealed normal architecture, with intact elastic lamellae, well-aligned smooth muscle cells, and preserved collagen distribution (data not shown).

**Figure 3.**
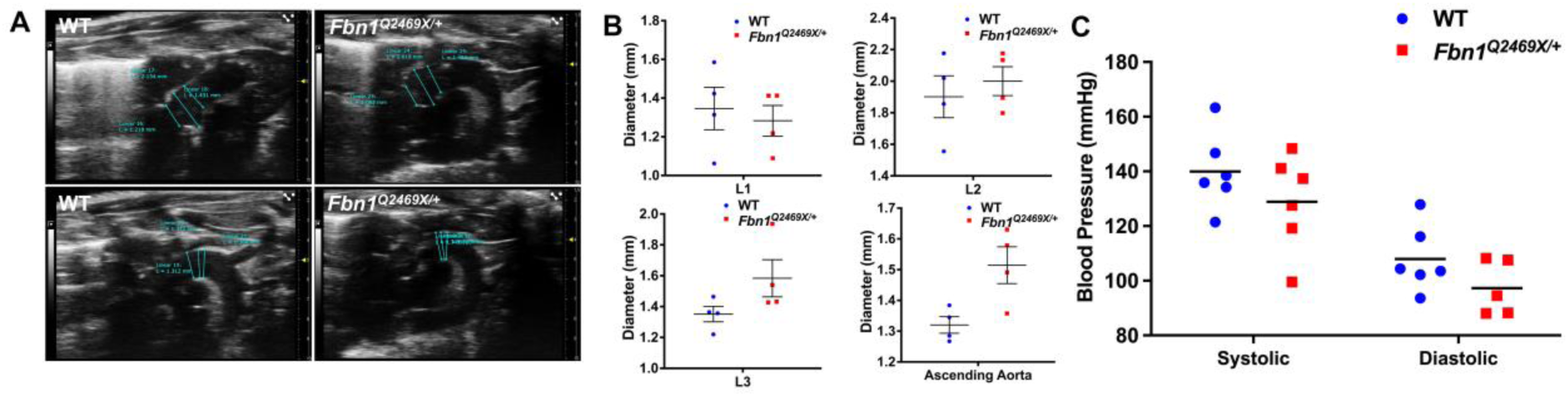
Morphometric and hemodynamic features of *Fbn1^Q2469X/+^*mice. (A) Representative echocardiographic images showing aortic dimensions at the annulus (L1), sinus of Valsalva (L2), and sinotubular junction (L3) in *Fbn1^Q2469X/**+**^* mice (right) and wild-type (WT) littermates (left). (B)Quantitative measurements of aortic diameters at L1–L3 and the ascending aorta. *Fbn1^Q2469X/**+**^* mice exhibit mild, non-significant increases in diameter (∼0.2 mm at L3 and ∼0.3 mm at the ascending aorta; n = 4 per group). (C) Systolic and diastolic blood pressure in in *Fbn1^Q2469X/**+**^* and WT mice show no significant differences. Data are presented as mean ± SEM; statistical analyses by unpaired t test.

### *Fbn1^Q2469X/Q2469X^*mice develop progressive thoracic aortic aneurysms, leading to sudden death due to aneurysm rupture

The *Fbn1^Q2469X/Q2469X^* mice all die out within 25 days after birth while the littermate *Fbn1^+/+^* and *Fbn1^Q2469X/+^* mice are alive during the time. To visualize aneurysm progression at different time after birth, we performed latex injection in *Fbn1^Q2469X/Q2469X^*mice (Figure 4A). We do not observe apparent difference among littermates with different genotypes from P0-P5 after birth. The dilation of the aortic root and ascending aorta dilation is observed at P7 after birth (Figure. 4A Early stage). The dilation progress to rupture is observed at P13 at the aortic root, ascending aorta-arch region (Figure 4A, late stage). The aneurysm rupture is observed at P13 at the junction of ascending aorta (Figure 4A, right side perfused by PBS).

**Figure 4.**
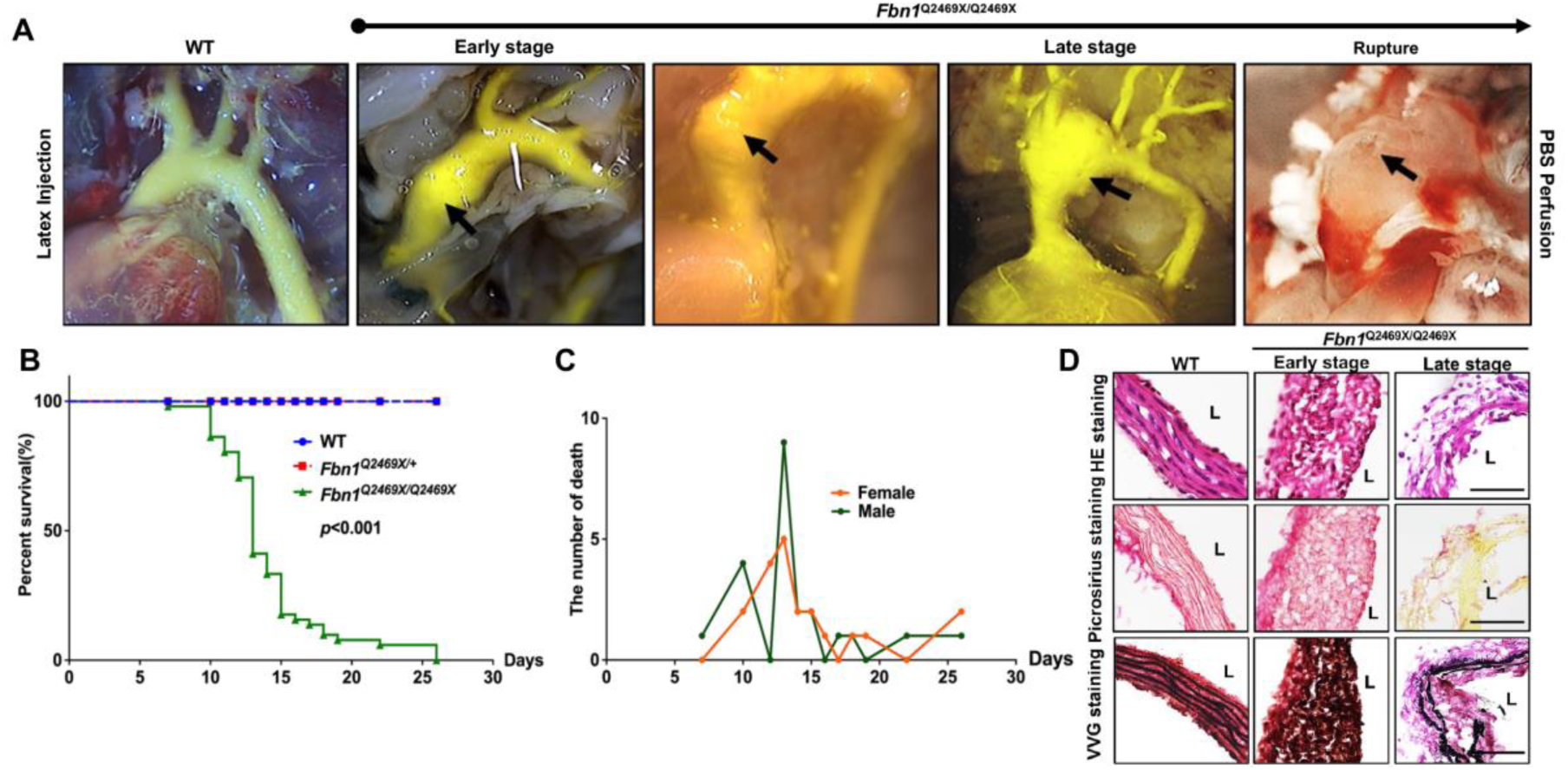
*Fbn1^Q2469X/Q2469X^* mice develop consistent progressive aneurysm, leading to rupture. (**A**) *Fbn1^Q2469X/Q2469X^* mice show rapid aorta dilation during aneurysm progression from early stage, late stage to rupture. The images demonstrate the aneurysm progression and rupture in aortic root, ascending aorta and arch by the perfusion with PBS and followed by latex injection. Black arrows indicate the location of aneurysm dilation and rupture. (**B)** The Kaplan-Meier survival curve of mice for wild type (n=51), *Fbn1^Q2469X/+^* (n=70) and *Fbn1^Q2469X/Q2469X^*(n=51) mice. *P*<0.001. (**C**) There is no difference between male and female *Fbn1^Q2469X/Q2469X^* mice ( n=20 for female; n=22 for male). (D) The pathological changes of aneurysm at ascending aorta during aneurysm progression from early stage to late stage. The H&E staining shows changes of the thickness, SMC organization in the vessel wall during aneurysm progression (Top). Picro sirius red staining shows collagen degradation in the aorta at late stage (middle). Verhoeff Van Gieson staining shows elastin disorganization at early stage and fragmentation at late stage.

To determine the mortality rate of *Fbn1^Q2469X/Q2469X^*mice, we performed Kaplan-Meier survival curve analyses of *Fbn1^+/+^*, *Fbn1^Q2469X/+^* and *Fbn1^Q2469X/Q2469X^*mice, and the log-rank test determines the statistical significance between *Fbn1^Q2469X/Q2469X^* mice and *Fbn1^+/+^*or *Fbn1^Q2469X/+^*mice (n=30 for each genotype, *P*<0.0001): all of the *Fbn1^Q2469X/Q2469X^* mice die of aneurysm rupture within 25 days, and most of these mice die between 13-15 days after birth (Figure 4B). All *Fbn1^Q2469X/Q2469X^*mice die from hemothorax with blood clots filled lung and the rupture of thoracic aortic aneurysm. Sex does not affect the survival of *Fbn1^Q2469X/Q2469X^*mice (Figure 4C).

To investigate the molecular changes of the vessel wall during aneurysm progression to rupture, we conducted histological analyses using HE stain, Picrosirius Red connective tissue stain and Verhoeff Van Gieson Elastin Stain on the thoracic aortic sections of the aneurysms at both early and late stages (Figure 4D). We observed that (1) the vessel wall of aneurysms is thickened at early stage and degenerated at late stage; (2) the smooth muscle layer of the aorta wall become disorganized in *Fbn1^Q2469X/Q2469X^* mice comparing with the elongated SMC arrays in the wild type mice; (3) the cell loss is apparent at the late stage; (4) Picrosirius Red connective tissue staining shows no significant changes in collagen in early-stage aneurysms, but drastic collagen degradation in late-stage aneurysms; (4) Verhoeff Van Gieson Elastin Stain detects amorphous elastic fibers at early stage and severe fragmentation and degradation at late stage in *Fbn1^Q2469X/Q2469X^* mice, not in the wild type littermates. These results demonstrate the pathological changes including VSMC dedifferentiation, collagen degradation and elastin fragmentation during aneurysm progression in *Fbn1^Q2469X/Q2469X^* mice during aneurysm progression.

### Inflammation and immune response are the predominant pathogenic processes in *Fbn1^Q2469X/Q2469X^* aneurysms

To explore the molecular drivers of aneurysm progression to rupture, we performed bulk RNA-seq analyses of thoracic aortas from *Fbn1^Q2469X/Q2469X^* mice and wild-type (WT) littermates (n=3 per group; Figure 5A). Hierarchical clustering revealed distinct transcriptomic patterns between genotypes under a threshold of *P*<0.01 (Figure 5B). Using established criteria (fold change >2.0, *P*<0.01, FDR<0.05), we identified 854 differentially expressed genes (DEGs), including 643 upregulated and 211 downregulated transcripts in *Fbn1^Q2469X/Q2469X^ mice*.

**Figure 5.**
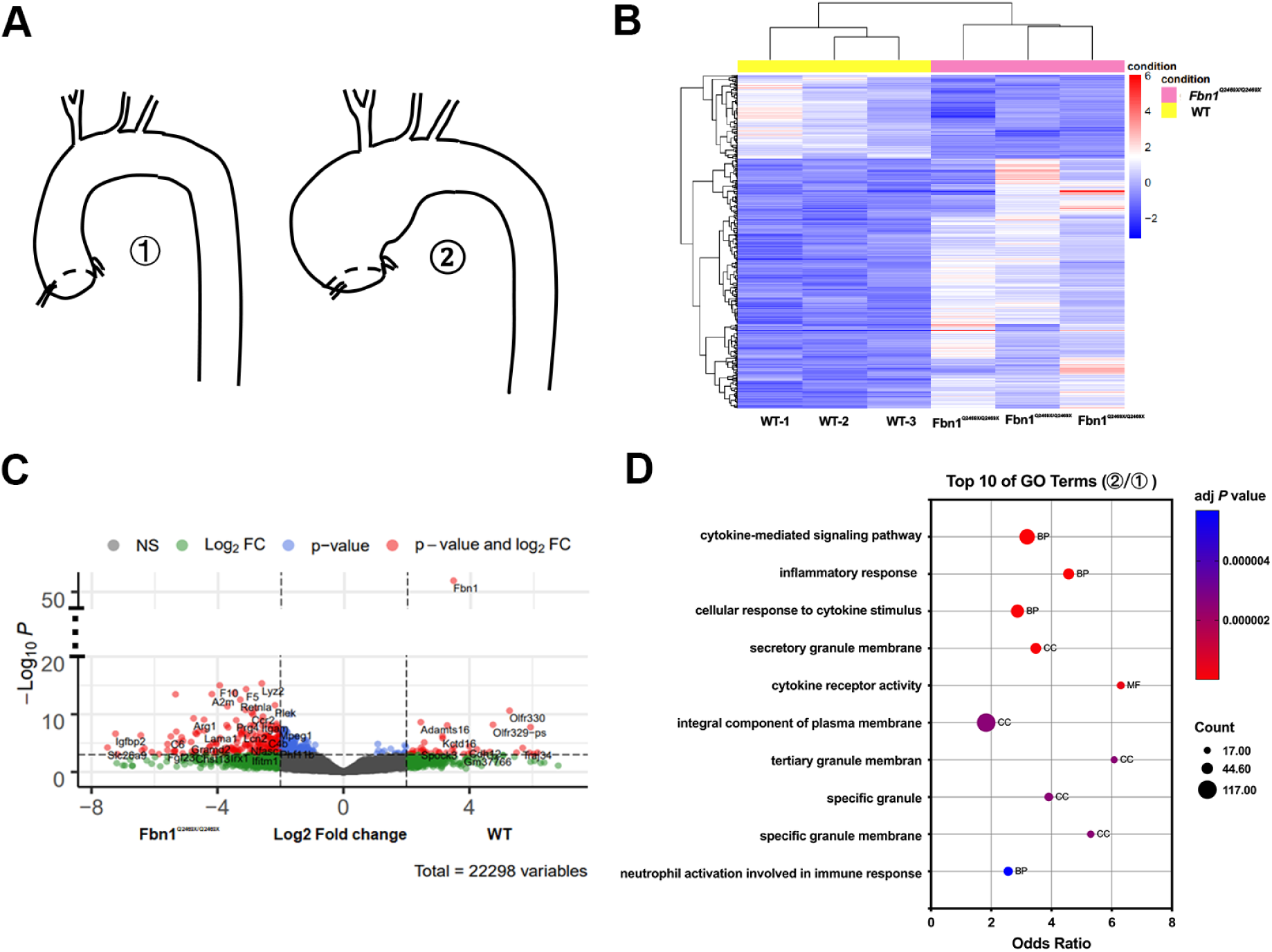
RNA-seq analyses reveal altered gene expression profiles in *Fbn1^Q2469X/Q2469X^* aneurysmal aortas. (A) Thoracic aortas from wild-type (WT) and *Fbn1^Q2469X/Q2469X^*mice (n=3 per group) were collected for RNA extraction and RNA-seq analysis. (B) Heatmap of the most variable genes (adjusted *P*<0.01), generated from variance-stabilized counts. The color scale reflects z-scores from the mean. Pearson correlation between samples was calculated using variance-stabilized counts, and sample distance was visualized as sqrt(1–cor²). (C) Volcano plot showing differentially expressed genes between WT and *Fbn1^Q2469X/Q2469X^* mice. The y-axis represents –log10 *P* values (*P*<0.01), and the x-axis represents log2 fold change. (D) Top 20 Gene Ontology (GO) terms enriched among differentially expressed genes, identified using DAVID. Red bars represent Biological Process terms, and the green bar represents a Molecular Function term.

A volcano plot highlighted significantly altered DEGs between mutant and WT aortas, including genes associated with immune response, thrombosis, vascular remodeling, and extracellular matrix turnover, such as *F5, F10, Arg1, A2M, Thbs1, Itgam, Igfbp2, C6, Retnla,* and *ADAMTS16* (Figure 5C). Gene Ontology (GO) enrichment analysis of DEGs identified 148 biological process, 10 molecular function, and 21 cellular component terms. The top 20 enriched GO terms (adjusted *P*<0.05) included cytokine-mediated signaling pathways and inflammatory responses as the most significantly altered categories during aneurysm progression and rupture (Figure 5D).

KEGG pathway analysis revealed enrichment of 29 pathways among upregulated DEGs and 3 pathways among downregulated DEGs (Figure S1). Cytokine–cytokine receptor interaction and primary immunodeficiency were the most significantly enriched pathways in upregulated and downregulated DEGs, respectively (adjusted *P*<0.05). Together, these transcriptomic analyses highlight inflammation and immune activation as the predominant pathogenic processes driving aneurysm progression in *Fbn1^Q2469X/Q2469X^* mice.

### Gene Ontology enrichment highlights inflammatory signaling in aneurysms of *Fbn1^Q2469X/Q2469X^* mice

To further define the biological processes underlying the transcriptomic changes observed in aneurysms of *Fbn1^Q2469X/Q2469X^* mice, we performed Gene Ontology (GO) enrichment analysis on the differentially expressed genes (DEGs) identified by RNA-seq. A chord diagram illustrated the top 10 enriched GO terms and their associated DEGs, with upregulated genes (red) and downregulated genes (purple) mapped to functional categories such as cytokine signaling, immune activation, and extracellular matrix organization (Figure 6A).

**Figure 6.**
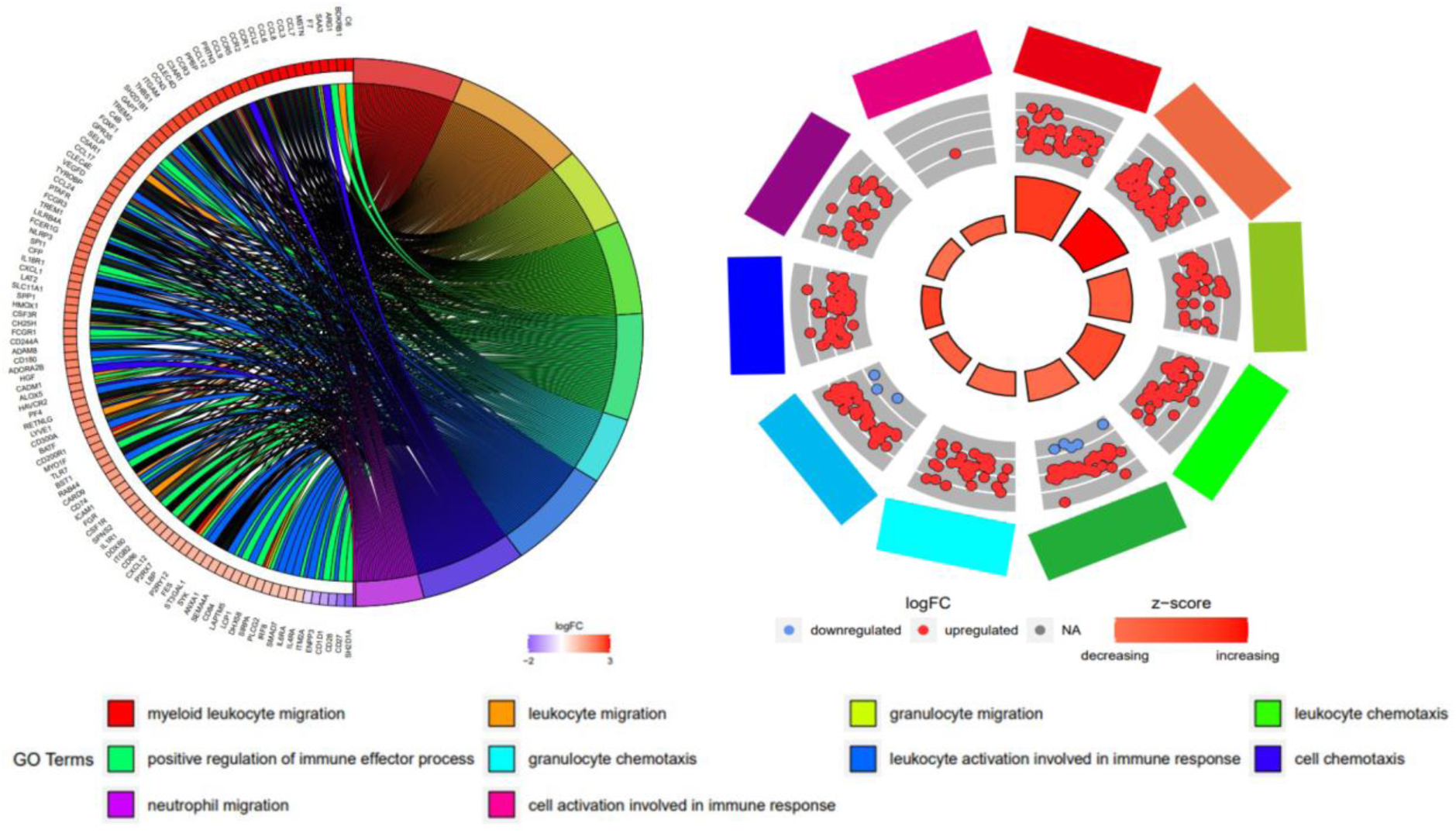
Gene Ontology enrichment analysis highlights inflammatory pathways in *Fbn1^Q2469X/Q2469X^* aneurysmal aortas. (A) Chord diagram showing the top 10 enriched GO terms and associated significantly differentially expressed genes between *Fbn1**^Q^***^2469^***^X/Q2469X^*** and WT aortas. Genes are color-coded by log2 fold change, with red indicating upregulation and purple indicating downregulation. GO terms are color-coded as indicated at the bottom. (B) Circular plot illustrating the top 10 inflammation-related GO terms. The outer ring represents individual GO terms, with upregulated genes in red and downregulated genes in blue. The inner circle is scaled by *P* value (larger size = greater significance) and color-coded by z-score, where red indicates pathway activation and blue indicates pathway suppression.

A circular plot analysis of the top 10 inflammation-related GO terms further emphasized immune signaling as a central feature of the transcriptomic profile (Figure 6B). Upregulated genes were prominently linked to pathways involved in cytokine-mediated signaling, leukocyte activation, and response to wounding. The z-score distribution indicated that most inflammatory pathways were activated in aneurysms of *Fbn1^Q2469X/Q2469X^*mice, with pathway significance reflected by circle size (*P*<0.05). Together, these analyses highlight inflammation as a dominant biological process in aneurysm progression, building upon the global transcriptomic analyses.

### Adventitial inflammation increases with aneurysm progression

To confirm the presence of immune cell infiltration marked by CD11B^+^ in aneurysms, we performed immunofluorescent staining on OCT sections from the thoracic aorta of *Fbn1^Q2469X/Q2469X^* mice at early, intermediate and late stages. Immunofluorescence staining reveals the spatial distribution of immune cells in the vessel wall: aneurysmal vs nonaneurysmal area; few immune cells are detected in the endothelial nor VSMCs, most immune cells are located in the adventitia area. These results demonstrated a progressive increase in adventitial immune cell infiltration as aneurysms advanced from early to late stages. At the early stage, few CD11B⁺ cells were observed in the aneurysmal vessel wall. By the intermediate stage, CD11B⁺ cells became localized primarily to the adventitia of the aneurysmal region. At the late stage, CD11B⁺ signals were markedly elevated, with dense accumulation particularly in the adventitia near rupture sites (Figure 7).

**Figure 7.**
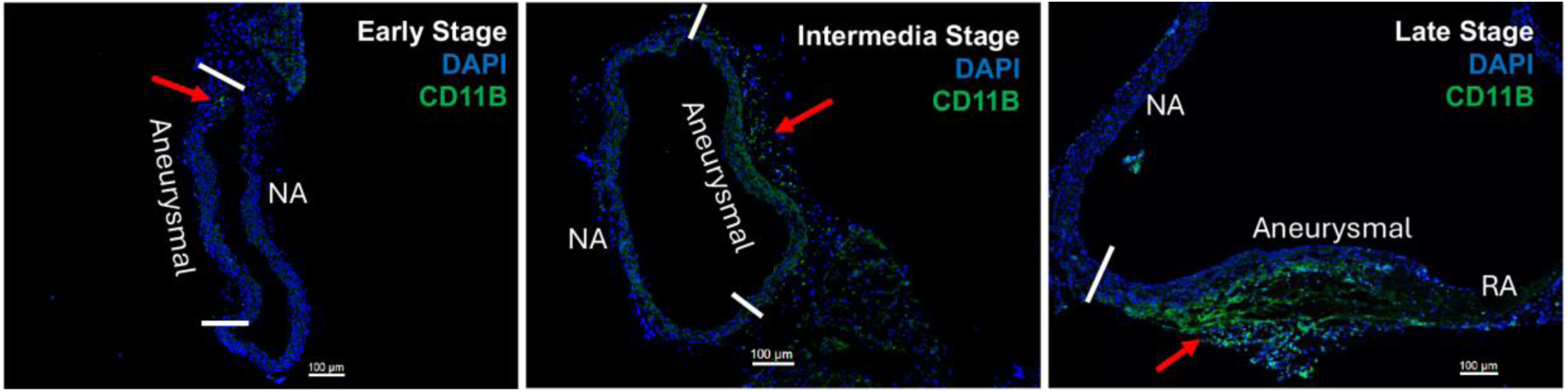
Progressive adventitial inflammation during aneurysm development. Immunofluorescence staining for CD11B demonstrates increasing immune cell infiltration in the adventitia of the aneurysmal vessel wall from early to late stages. At the early stage, few CD11B⁺ cells are detected in the aneurysmal wall. At the intermediate stage, CD11B⁺ cells are localized primarily to the adventitia of the aneurysmal region. At the late stage, CD11B⁺ signals are markedly increased in the adventitia aneurysmal area near the rupture area. NA, nonaneurysmal area; RA, rupture area.

This stepwise increase in adventitial immune cell infiltration suggests that inflammation in the outer vessel wall is closely associated with the pathological remodeling that drives aneurysm progression. The pattern of CD11B⁺ cell accumulation highlights the adventitia as a critical site of immune activation in thoracic aortic aneurysms.

## Discussion

### Summary of key findings

In this study, we generated a *Fbn1*^Q2469X/+^ mouse model carrying a patient-derived nonsense mutation that results in *Fbn1* deficiency due to nonsense-mediated decay, rather than production of a truncated protein (Figure 1, Figure 2). Heterozygous mice appear normal with only mild, non-progressive changes in aorta dimensions (Figure 3), whereas homozygous mice develop rapidly progressive thoracic aortic aneurysms that proceed through reproducible stages and culminate in rupture between 10 and 25 days of age (Figure 4). Histological analyses revealed smooth muscle cell remodeling, collagen degradation, and elastic fiber fragmentation (Figure 4). Bulk RNA-seq and immunostaining analyses reveal that inflammation becomes highly enriched at late stages (Figure 5-7), suggesting that immune activation is a critical component of terminal aneurysm progression. Together, these findings establish an aneurysm mouse model that provides a valuable platform for mechanistic studies and therapeutic discovery (Figure 8).

**Figure 8.**
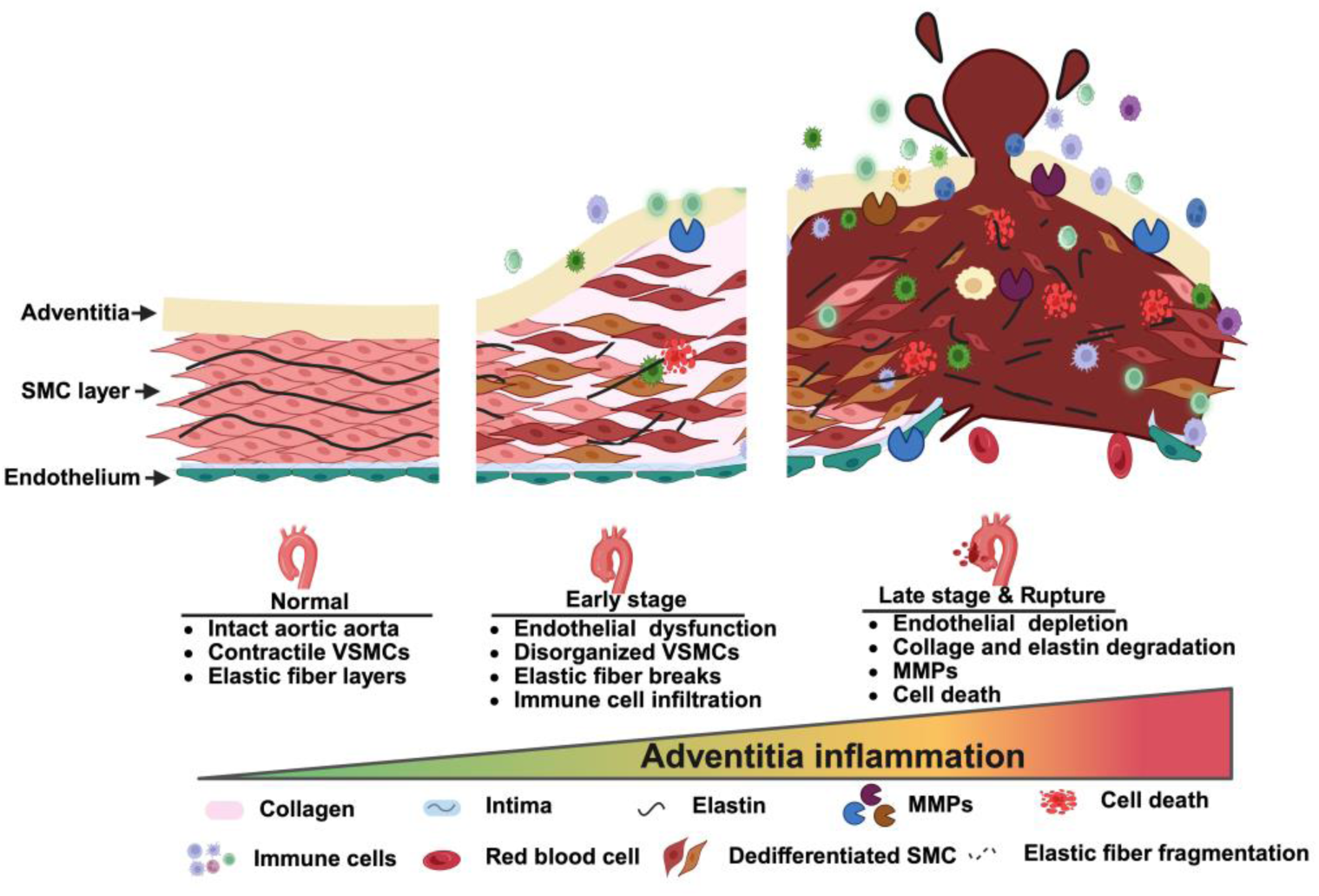
Summary of aneurysm progression and adventitial inflammation in FBN1Q2469X mouse model. Schematic model illustrating the structural and molecular changes during aneurysm progression from normal aorta to rupture. In the normal vessel, the aortic wall consists of intact endothelium, organized smooth muscle cells (SMCs), and elastic fibers. At early stage, aortic dilation is accompanied by SMC disorganization, elastic fiber breaks, and initial immune cell infiltration. In late-stage aneurysm and rupture, the vessel wall exhibits marked adventitial inflammation, collagen and elastic fiber degradation, matrix metalloproteinase (MMP) activation, cell death, and endothelial depletion. The gradient highlights the progressive increase of adventitial inflammation as a central driver of aneurysm progression to rupture. Symbols denote histological and molecular features, including collagen, elastin, intima, MMPs, immune cells, dedifferentiated SMCs, and elastic fragmentation.

### Inflammation in aneurysm pathogenesis as a late-stage driver

The contribution of inflammation to Marfan-associated aortic aneurysms has been debated due to early studies reporting minimal immune activation involvement^29,30^. Recent single-cell RNA sequencing analyses of aortic tissue from Marfan patients and Marfan mouse models have, however, revealed extensive immune cell infiltration and cytokine signaling^31,32^, suggesting that inflammation contributes to disease progression. Consistent with these observations, our transcriptomic and histological analyses show that strong enrichment of cytokine-mediated signaling, leukocyte activation, and immune pathways during the late phase of aneurysm development in the *Fbn1^Q2469X/Q2469X^* model. Immunofluorescence assays demonstrated progressive accumulation of CD11B⁺ immune cells in the adventitia near the ruptured region. These results indicate that adventitial inflammation becomes a prominent feature of only during the late phase of aneurysm development and is not an early initiating event.

### Aneurysm morphology and relevance to Marfan syndrome

A concern regarding the *Fbn1^+/Q2469X^* model is that late-stage saccular aneurysms differ from the fusiform morphology typically seen in Marfan syndrome, where the root and ascending aorta are most commonly affected. In our mouse model, aneurysms are fusiform at the early stage and evolve into a saccular morphology at the late stage (Figure 4A). Clinically, Marfan aneurysms undergoing elective surgery are typically repaired before reaching advanced pathological stages, which may explain why late-stage saccular remodeling is less often documented in patients. Although saccular aneurysms are rare, they have been reported in individuals with fibrillin-1–related connective tissue disorders disorders^28^ (https://thoracickey.com/marfan-syndrome-aortic-aneurysm-and-dissection/). These isolated observations suggest that aneurysm morphology can vary among individuals and across different stages of disease progression. Thus, while not an exact phenocopy of classical Marfan aortopathy, the late-stage saccular morphology in *Fbn1^Q2469X/Q2469X^* mice should be viewed as an experimental manifestation that still captures key pathogenic processes associated with fibrillin-1 deficiency.

### Comparison of the *Fbn1^Q2469X/+^* model with commonly used *Fbn1* mouse models

Existing two widely used Marfan mouse models (*Fbn1^C1041G/+^* and *Fbn1^mgR/mgR^*) have substantially advanced our understanding of aneurysm formation^25,26^, but each has inherent limitations. The *Fbn1^C1041G/+^* Marfan mouse model carries a patient-derived missense mutation and exhibits aorta dilation at the root and ascending aorta without dissection nor rupture^19^, limiting its utility for studying late-stage aneurysm pathogenesis. The *Fbn1^mgR/mgR^* Marfan mouse model was generated by an ES-cell recombination event that reduces *Fbn1* expression^20^, producing a hypomorphic state rather than a clinically validated patient mutation. Similar to several heterozygous *Fbn1* models including *Fbn1^mgR/+^* and *Fbn1^Δ/+^*, the heterozygous *Fbn1^Q2469X/+^*mice remain largely unaffected under normal conditions. The *Fbn1^mgR/mgR^* model, although widely used, exhibits significant variability in the timing and severity of aneurysm progression, making stage-specific analyses difficult to manage. Conversely, complete *Fbn1* knockout mice *Fbn1^Δ/Δ^*die shortly after birth^27^, preventing the study of postnatal aortic pathology.

Our homozygous *Fbn1^Q2469X/Q2469X^* mice also have a short lifespan, but these homozygotes retain ∼10% residual *Fbn1* transcript due to incomplete degradation through NMD (Figure 2C), producing reproducible, staged aneurysm progression that consistently culminates in rupture (Figure 4). Taking together, although *Fbn1^Q2469X/Q2469X^* model does not capture the full spectrum of Marfan-associated aortic phenotypes, it provides a consistent system for studying staged aneurysm progression.

### Translational relevance and conclusion

Consistent with our findings (Figures 5–6), recent transcriptomic and molecular analyses of human Marfan aneurysms and related mouse models have revealed overlapping pathogenic processes involving inflammation, oxidative stress, metabolic dysregulation, mechanotransduction defect, and mitochondrial impairment^6,31–33^. The involvement of nonsense-mediated mRNA decay (NMD) in *FBN1* haploinsufficiency underscores the relevance of patient-derived loss-of-function mutations in shaping disease biology^34^. Although The *Fbn1^+/Q2469X^* mouse model does not reproduce the full spectrum of Marfan syndrome, it offers a genetically defined system in which late-stage aneurysm progression can be examined with consistency. By linking a clinically relevant *FBN1* mutation to aneurysm pathology, this model enables mechanistic investigation of extracellular matrix–based drivers of aneurysm progression and supports the evaluation of therapeutic strategies aimed at stabilizing the aortic wall in aortic aneurysm diseases.

## Acknowledgments

This project was initiated upon the suggestion and encouragement of Dr. Lynne Maquat (University of Rochester). Her insights into nonsense-mediated decay were instrumental in our pursuing the generation of a Marfan mouse model carrying a FBN1 nonsense mutation. We would like to thank the excellent technique assistance from people working in the lab and help discussions from colleagues during the course of this study. We are grateful for the aneurysm tissues of Marfan patient from the Biobank of University of Michigan. This work was supported by funding from the WSU OVPR Grant Boost Award, the National Heart, Lung, and Blood Institute (HL119815, HL142956 to L.L.).

## Author contributions

S.W., J.Z. A.P. and L.L. conceived and designed the experiments; S.W., J.Z. A.P, D. J., M.Y., performed the experiments; S.W., J.Z. A.P, D. J., S.Y., M.Y., D.X., C.L., L.L., Y.X., K.Z., H.L., Z.Y. and L.L. analyzed the data; S.W., J.Z. A.P., M.D., B.Y., K.Z.,Y.X. and L.L. wrote and revised the manuscript. All authors approved the manuscript.

## Additional information

The authors declare no competing financial and non-financial interests.

**Figure S1.**
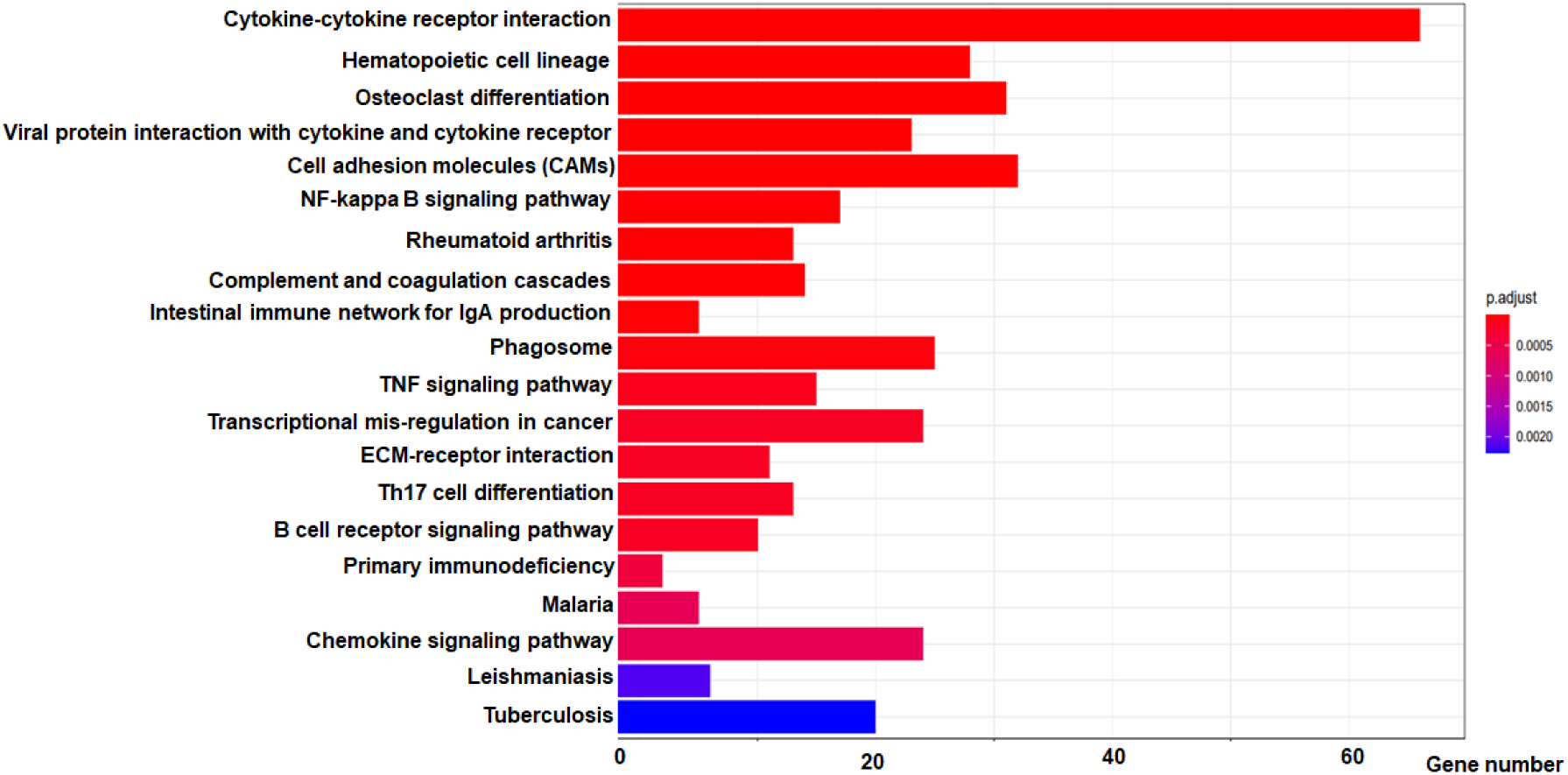
The KEGG analysis of RNA-seq data. The KEGG pathway enrichment analysis was generated with the significantly differentially expressed genes (*P*<0.01) in the RNA-seq of FBN1Q2469X/Q2469X mice and WT. Top 20 KEGG pathways are shown with bar plot: the color represent the adjusted p value of enrichment the KEGG pathways; X axis represents the gene number involved in the KEGG pathways.

**Table S1.** *FBN1* is highly conserved between human and mouse.

| ***FBN1*** | **Human** | **Mouse** |
| --- | --- | --- |
| **Chromosome** | **15** | **2** |
| **Gene size** | **237.4kb, 66 exons** | **66exons** |
| **Exons/ATG/TAA** | **66/2/66** | **66/2/66** |
| **c.7399C>T**<br>**transition** | **In Exon 59**<br>**(CAG-TAG)** | **In Exon 60**<br>**(CAG-TAG)** |
| **mRNA (NCBI)** | **11756bp** | **9847bp** |
| **Protein** | **2873aa (350KD)** | **2873aa** |
| **Protein identity** | **2765/2873= 96.24%** |  |
| **Protein similarity** | **2834/2873= 98.64%** |  |
| **a nonsense**<br>**mutation** | **Q2467X** | **Q2469X** |

